# Echinocandin tolerance and persistence *in vitro* are regulated by calcineurin signaling in *Candida glabrata*

**DOI:** 10.1101/2025.08.19.671059

**Authors:** Abigail A. Harrington, Timothy J. Nickels, Kyle W. Cunningham

## Abstract

Upon exposure to echinocandins, growing yeast cells begin to accumulate cell wall damage and eventually die, resulting in therapeutic effects. While resistance to echinocandins is well studied, tolerance and persistence mechanisms that may also contribute to clinical failures and relapses remain understudied. In time-kill assays with micafungin *in vitro*, the opportunistic pathogen *Candida glabrata* exhibited biphasic kinetics of cell death. Modeling with exponential decay equations distinguished a fast-dying major population from a slow-dying minor population, indicative of persistence. A genome-wide forward-genetic screen revealed dozens of genes that appeared to regulate persistence and/or tolerance, but not resistance. Several of those genes encoded calcineurin and its upstream regulators. Using individual gene knockout mutants and FK506, we show that calcineurin signaling increases the lifespans of most *C. glabrata* cells through a process that is largely independent of Crz1, one of its downstream effectors. The formation of long-lived persister-like cells (i.e. persistence) was strongly dependent on calcineurin signaling, independent of Crz1. Pre-activation of calcineurin using genetic or chemical stressors, such as manogepix, strongly increased tolerance and persistence in *C. glabrata*, suggesting antagonism of echinocandin efficacy by this new antifungal. Calcineurin signaling was also necessary for induction of tolerance and persistence in *Candida albicans*. The findings suggest that short-term administration of FK506 during the earliest stages of echinocandin treatment may improve clinical outcomes while possibly avoiding long-term immunosuppression.

**IMPORTANCE:** Treatment of fungal infections is often unsuccessful. Potential causes of antifungal failure include tolerance and persistence, which are poorly understood processes used by fungal pathogens to survive our assaults. This study utilizes detailed experimental protocols and genome-wide screens to discover how *Candida glabrata* induces tolerance and persistence to a major class of antifungals. The findings suggest that a clinical immunosuppressant may be repurposed to combat tolerance and persistence in this pathogenic yeast as well as *Candida albicans* and perhaps others.

## INTRODUCTION

At least four distinct processes contribute to antibiotic treatment failure: resistance, heteroresistance, tolerance, and persistence (1–3). Resistance occurs when microbes acquire mutations that alter the expression or function of the drug target, the influx, efflux, sequestration or metabolism of the compound, or otherwise increases the effective dosage of the antimicrobial drug. Common measures of antibiotic resistance include the minimal inhibitory concentration (MIC) or the concentration causing a 50% inhibition of maximal growth (IC50). Resistance mutations are stably inherited by daughter cells and cause infections that require higher doses of the antibiotic or alternative antibiotics to remedy. Heteroresistance occurs when a small subpopulation of cells within a clonal population acquires phenotypic resistance without any stable mutations. Heteroresistant subpopulations have a greatly increased MIC relative to the majority of cells in the population and have been shown to contribute to treatment failure in murine models of infection. Tolerance occurs when the clonal population has acquired mutations that increase the lifespan of the cells, even when exposed to excess antibiotics, without altering the MIC or IC50. To overcome tolerance, the duration of antibiotic treatment must be increased, not the dose. Tolerance is usually assayed using time-kill experiments that measure the number of “viable” cells in the population (colony forming units, or CFU) after transient exposure to supra-MIC doses of antibiotics and estimating the half-life of the population. Persistence refers to an epigenetic regulatory process that produces a small number of highly tolerant “persister cells” amongst the large population of fast-dying susceptible cells. Persister cells are typically quiescent, at least transiently, and are therefore not killable by multiple classes of antibiotics, which can increase the likelihood of relapse. All these processes have been studied thoroughly in many infectious species of bacteria.

Antifungal resistance is well-studied in some pathogenic fungi (4, 5), but heteroresistance, tolerance, and persistence are just beginning to be distinguished and individually unraveled at the molecular level (6). Tolerance and heteroresistance to azoles, fungistats targeting ergosterol biosynthesis in the ER, is now being investigated in several pathogenic species of yeasts (7–11) as there is growing evidence of clinical relevance of these processes (12). Heteroresistance to echinocandins, fungicides targeting the cell wall, was recently associated with breakthrough infections by *Candida parapsilosis* (13). A retrospective study found that rare echinocandin-tolerant strains of *Candida tropicalis* were much more lethal than non-tolerant strains in patients treated for candidemia (14). Though the molecular mechanisms of echinocandin heteroresistance and tolerance have not been elucidated, an inhibitor of the Ca^2+^/calmodulin-dependent protein phosphatase calcineurin (FK506) abolished the tolerance of *C. tropicalis* and substantially increased the lifespan of infected animals undergoing echinocandin treatment (14). A better understanding of the regulatory mechanisms responsible for heteroresistance, tolerance, and persistence could potentially lead to development of antifungal therapies that lack undesirable side-effects on the patient (such as immunosuppression caused by FK506).

*Candida glabrata* (also known as *Nakaseomyces glabratus*) is the second most common cause of life-threatening candidemia and candidiasis next to *C. albicans* (15–17), though it belongs to a genus that is more closely related to *Saccharomyces* than true *Candida* (18). Due to its innate and easily acquired resistance to antifungals and its rising incidence, the World Health Organization has classified *C. glabrata* as a High Priority Threat (19). In *C. glabrata*, azole resistance mutations arise primarily in the target (*ERG11*) and in a transcription factor gene (*PDR1*) that regulates expression of transporters that efflux the drug (20). *PDR1* also confers mild resistance to most echinocandins through expression of lipid flippases (21, 22). However, strong resistance to echinocandins arises primarily through mutations in the two targets, encoded by *FKS1* and *FKS2* (23, 24), the latter of which depends on calcineurin and the Crz1 transcription factor for maximal expression (25). When a resistance mutation arises in *FKS2*, its impact *in vitro* can be blocked by calcineurin inhibitors (25, 26). Even in the absence of antifungals, calcineurin also promotes virulence of *C. glabrata* in mouse models of invasive candidiasis through a mechanism that is potentially independent of Crz1 (27, 28). Time-kill experiments *in vitro* have shown most *C. glabrata* cells die quickly when exposed to high doses of echinocandins while a small and variable sub-population dies more slowly (29, 30), suggesting the possible development of long-lived persister cells. Such persister cells serve as a reservoir for acquisition of resistance mutations (30). After engulfment into phagosomes by macrophages, *C. glabrata* cells typically remain viable and survive longer when exposed to echinocandins, with an increased number of long-lived cells (31). Therefore, tolerance and persistence in this species may be clinically important and regulated by unknown mechanisms.

This study quantifies tolerance and persistence in *C. glabrata* exposed to echinocandins *in vitro* by fitting high resolution time-kill data to exponential decay equations developed previously for antibiotic research (2). It also implements a genome-wide genetic screen using Tn-seq to identify specific regulators of tolerance and persistence, as well as individual gene knockout experiments. The findings suggest tolerance and persistence are governed by processes distinct from resistance and heteroresistance. Remarkably, *C. glabrata*, and *C. albicans*, appeared to induce tolerance and persistence ‘on demand’ through the activation of a calcineurin. Therefore, FK506 and other drugs that block calcineurin signaling in yeasts and other fungi may improve clinical outcomes by lowering tolerance and persister cell development in addition to lowering resistance (expression of echinocandin targets). Conversely, the findings suggest that drugs, mutations, and host conditions that stress fungal cells and pre-activate calcineurin may promote tolerance and persistence, thereby worsening clinical outcomes.

## RESULTS

### Quantitation of echinocandin tolerance and persistence in *C. glabrata*

Time-kill assays have been used routinely to discriminate, quantify, and characterize sub-populations of cells that exhibit distinct rates of killing by antibiotics (2). These assays involve treatment of clonal populations with high doses of cidal drugs for varying lengths of time, plating the treated cells onto drug-free agar media at appropriate dilutions, and then counting the number of colonies that appear after additional incubation. The colony forming units (CFU) per mL of starter culture is then charted over time and, if biphasic, analyzed by fitting to the sum of two exponential decay equations (2). When 8 independent stationary-phase cultures of wild-type *C. glabrata* (strain BG14) were diluted into fresh medium containing 125 ng/mL micafungin, biphasic survival kinetics were observed after a brief lag (Fig. 1). When charted individually, the eight cultures all behaved similarly (Fig. S1). The lag disappeared when log-phase cultures were tested, while the biphasic kinetics remained (Fig S2). After excluding the lag and fitting the averaged CFUs to exponential equations (Fig. 1, smooth curve), a minor population (9%) of long-lived cells (t ½ = 3.75 hr) could be distinguished from the major population of fast-dying cells (t ½ = 0.75 hr). To test whether stable mutants contributed to the slow-dying sub-populations, eight colonies that arose after 24 hr treatments with micafungin were picked, regrown in fresh medium, and retested individually. All eight cultures again produced similar numbers of long-lived cells (Fig. 1, open symbols and dashed curve). These findings show that clonal cultures of *C. glabrata* consistently and transiently produce two phenotypically distinct populations of cells, the minor one with nearly 5-fold increased half-life in micafungin.

**Figure 1.**
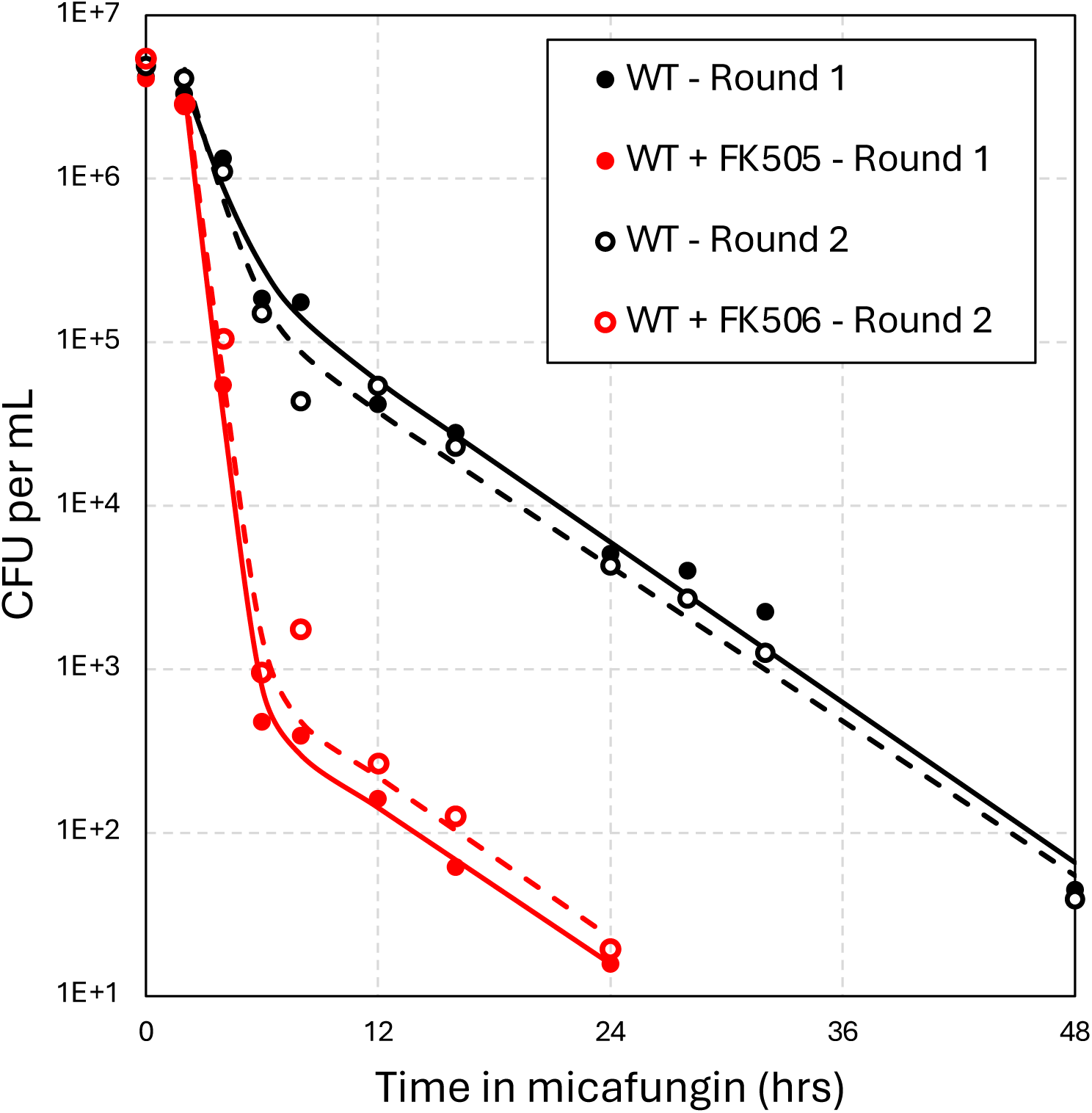
Quantifying tolerance and persistence in *C. glabrata* in response to micafungin treatment. Eight single colonies of wild-type strain BG14 were grown to saturation for 72 hours, then diluted 50-fold into fresh SCD medium containing 125 ng/mL micafungin alone (black) or micafungin plus 1 µg/mL FK506 (red). Cultures were shaken at 30°C, sampled at various timepoints, serially diluted, and plated on drug-free YPD medium. Colony forming units (CFU) were determined and the eight replicates averaged and charted (filled symbols). The best fit of the data to exponential decay equations are shown (solid smooth curves). Eight single colonies that appeared after 24 hours of treatment were picked and retested in the same conditions (Round 2; open symbols and dashed curves).

When micafungin was substituted with other glucan synthase inhibitors (e.g. caspofungin, ibrexafungerp), similar numbers of persister cells with similar lifespans were detected, though slight variation was evident (Fig. S3). The half-lives and the population sizes did not change even when the doses of these antifungals were greatly increased (Fig. S3). Additionally, the survival kinetics of several mutants that decreased (*fks2Δ*, *pdr1Δ*) or increased (*mrp20Δ*) resistance to micafungin (21) were indistinguishable from those of the BG14 parent strain (Fig. S4). Altogether, these results suggest *C. glabrata* can produce substantial numbers of long-lived persister-like cells through transient regulatory processes that are distinct from stable mechanisms that control resistance. Further, this mechanism also occurs independent of heteroresistance, as it was not detectable in wild-type *C. glabrata* strains (7). In the remainder of this study, the process of producing and maintaining such persister-like cells will be termed ‘persistence’ while the processes governing lifespan of the remaining susceptible cells will be referred to as ‘tolerance’, in accordance with earlier conventions (2).

### A forward genetic screen identifies regulators of tolerance and persistence

To identify genes that specifically regulate tolerance and/or persistence in *C. glabrata*, a genome-wide Tn-seq screen was implemented using a complex pool of transposon insertion mutants in strain BG14 that had been previously analyzed for resistance to micafungin and other antifungals (21, 32). Resistance screens employed very low doses of antifungals and long exposure times (48 hr), which would not have uncovered the genes that regulate tolerance and persistence. To elucidate such genes, the transposon pool was exposed to a high, lethal dose of micafungin for a short duration (6 hr) and then the cultures were washed and regrown in drug-free fresh medium before Tn-seq analysis. Mutants with elevated tolerance or persistence would be enriched in this modified protocol (i.e. positive Z-scores) while mutants with diminished tolerance or persistence would be depleted from the general pool (i.e. negative Z-scores).

Tolerance/persistence Z-scores were calculated for 5275 annotated genes by comparing the 6 hr and 0 hr exposures to 64 ng/mL micafungin (Table S1). When these Z-scores were charted against resistance Z-scores obtained previously with 8 ng/mL micafungin (21), the correlation was poor (Fig. 2). Hundreds of mitochondrial genes (Fig. 2, yellow symbols) and other genes that significantly increased resistance to micafungin had little impact on tolerance/persistence, as expected from the time-kill analysis of mitochondrial *mrp20Δ* mutants (Fig. S4). Conversely, dozens of genes with significantly high or low tolerance/persistence Z-scores had resistance Z-scores close to zero (Fig. 2). These findings further suggest that micafungin tolerance and persistence may be controlled by a relatively small number of genes that are largely distinct from those that regulate micafungin resistance.

**Figure 2.**
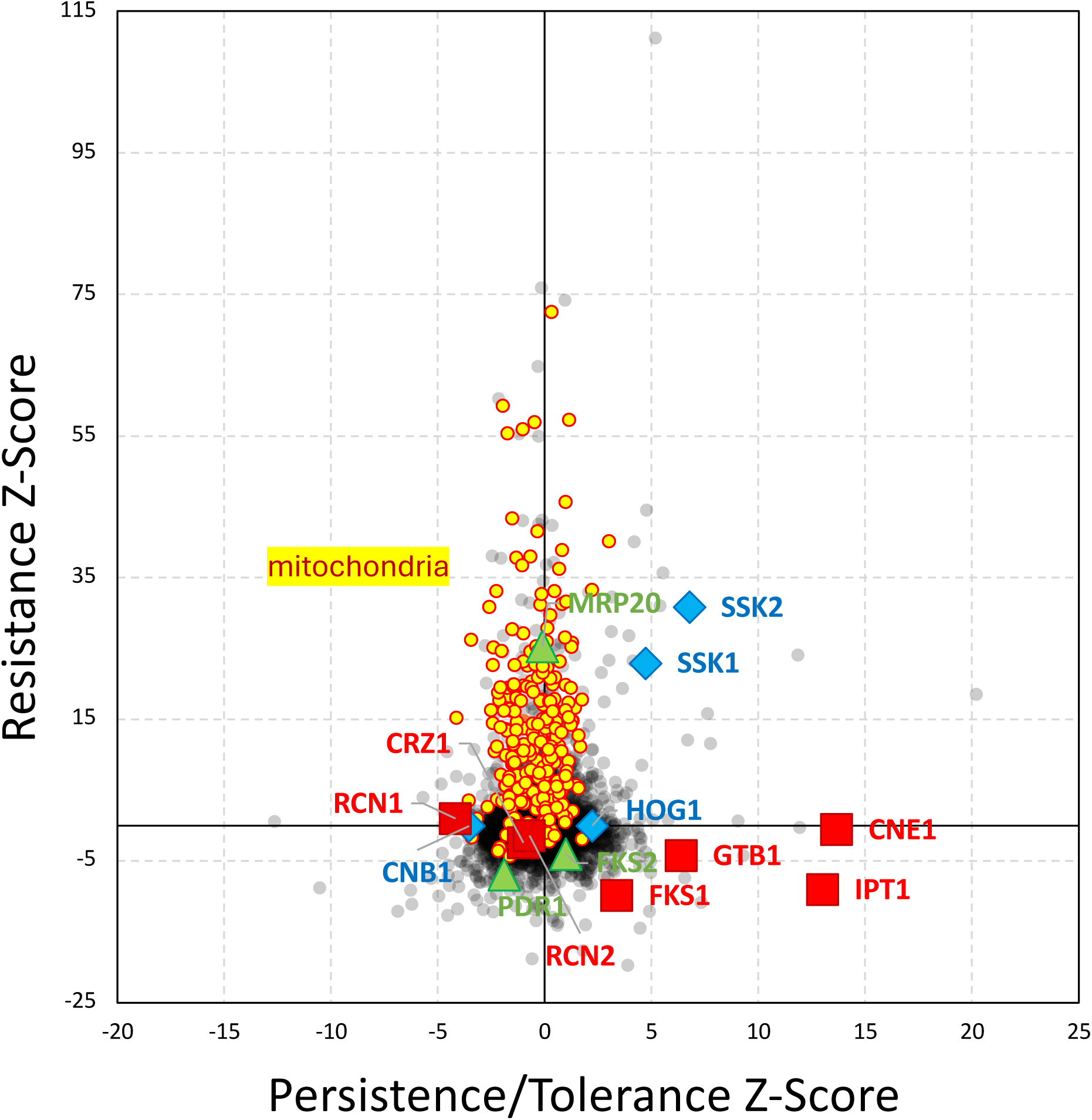
Genetic regulation of tolerance/persistence differs from that of resistance. A pool of Hermes transposon insertion mutants in the BG14 strain that was previously analyzed for resistance to low micafungin (Resistance Z-score) was reanalyzed in high micafungin for tolerance and persistence to high micafungin (Persistence/Tolerance Z-scores) and charted.

A total of 36 genes exhibited diminished tolerance/persistence (Z < −3.0) without exhibiting diminished resistance (Z > −3). GO term analysis (33) indicated significant enrichment (p-value = 5.2E-5; false discovery rate = 6.3E-2) of two small genes (*CNB1, RCN1*) that encode regulators of calcineurin, a well-studied Ca^2+^/calmodulin-dependent protein phosphatase. An upstream activator of calcineurin in *S. cerevisiae* and *C. albicans* (*KCH1*) (34, 35) also was present on this list. However, the *CNA1* gene, encoding the catalytic subunit of calcineurin, was not significant (Z = 0.26) possibly due to the small effective size of this gene and the presence of an autoinhibitory domain at the C-terminus. To explore the possible involvement of calcineurin in the regulation of tolerance and persistence, micafungin time-kill experiments were performed on wild-type *C. glabrata* in the presence of FK506, a specific inhibitor of calcineurin. Strikingly, FK506 caused a 2.5-fold decrease in the half-life of the susceptible cells (i.e. tolerance) and a 270-fold decrease in the number of long-lived persister-like cells in micafungin (Fig. 1, red symbols and curves). Of eight cells that survived 24 hr exposure to micafungin plus FK506 and produced colonies in drug-free media, all produced wild-type patterns in time-kill experiments upon retesting in the same conditions (Fig. 1 dashed curve, Fig. S1). These findings using FK506 confirm the Tn-seq results, suggest that calcineurin signaling drives both tolerance and persistence, and demonstrate those behaviors are readily reversible in *C. glabrata*. As calcineurin signaling becomes activated in response to micafungin and other stressors of the cell wall (36), tolerance and persistence may be protective behaviors in *C. glabrata* that are induced in response to stresses.

### Calcineurin drives tolerance and persistence in *C. glabrata* independent of Crz1 and Rcn2

The *CNA1* and *CNB1* genes were each knocked out in the BG14 parent strain and analyzed by time-kill experiments and mathematical modeling. Both the *cna1Δ* and *cnb1Δ* mutants exhibited strongly diminished tolerance and persistence relative to the wild-type control (Fig. 3, left panel). Though the number of persister-like cells decreased by more than 100-fold in both mutants relative to wild-type, the half-lives of the persister cells remained constant in all strains at approximately 3.2 hr. As expected, the presence of FK506 in the time-kill experiments did not impact the behavior of *cna1Δ* and *cnb1Δ* mutants and forced the wild-type parent strain to behave like the calcineurin-deficient mutants (Fig. 3, right). These findings further suggest that calcineurin signaling drives both tolerance and persistence upon exposure to micafungin.

**Figure 3.**
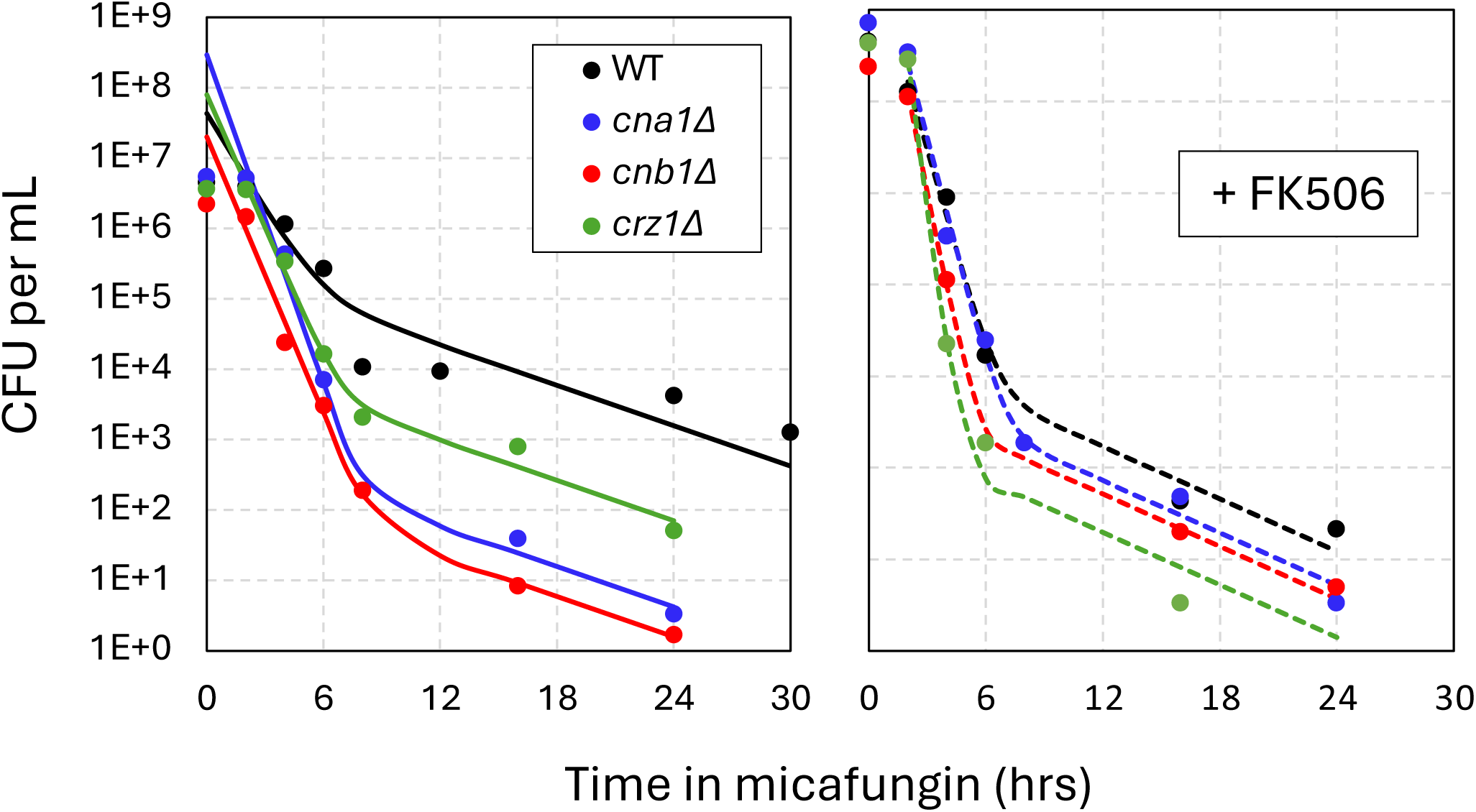
Calcineurin promotes tolerance and persistence. Time-kill assays of *cna1Δ* (blue), *cnb1Δ* (red), *crz1Δ* (green), and wild-type parent strain (black) were performed as described Fig 1. The averages of four biological replicates (symbols) and were fit to exponential decay equations (smooth curves). Experiments containing FK506 (1 µg/mL) were performed in parallel (right panel).

An important effector of calcineurin signaling is the transcription factor Crz1 (27). Though the *CRZ1* gene was not significant the Tn-seq screen, a *crz1Δ* mutant exhibited levels of tolerance and persistence that were intermediate between wild-type and the *cna1Δ* and *cnb1Δ* mutants (Fig. 3, left). In the presence of FK506, the *crz1Δ* mutant resembled the calcineurin-deficient strains (Fig. 3, right). Thus, Crz1 seemed to be required for a portion of the effects of calcineurin. Targets of Crz1 include *FKS2* and *RCN2* (25, 28, 37) neither of which were significant in the Tn-seq screen. An *fks2Δ* mutant was indistinguishable from wild-type in time-kill experiments (Fig. S4), suggesting it is not required for calcineurin-induced tolerance and persistence. An *rcn2Δ* mutant exhibited wild-type levels of tolerance and persister-like cells, but interestingly the half-life of the persister-like cells increased by ∼1.5-fold (Fig. S5). FK506 decreased the tolerance and persistence of *rcn2Δ* mutants to the same levels as the calcineurin-deficient mutants (Fig. S5). A plasmid that overexpresses *RCN2* from a strong constitutive *PDC1* promoter did not alter micafungin tolerance or persistence relative to a control plasmid when introduced into *cna1Δ* mutants or wild-type cells (Fig. S5). These findings suggest that Rcn2 functions in its canonical role as a feedback inhibitor of calcineurin signaling (28, 37) instead of an effector of calcineurin signaling in the regulation of tolerance and persistence.

To further explore the role of calcineurin in regulating tolerance and persistence, the effects of manogepix were studied. Manogepix is a preclinical antifungal that blocks GPI anchor biosynthesis in the ER (38) and produces cellular stresses that very strongly activates calcineurin signaling in *C. glabrata* (36). A 2-hour pre-treatment of wild-type cells with manogepix resulted in moderately increased tolerance and persistence to micafungin (Fig. 4). Though no such increases were observed in *cnb1Δ* mutants, the *crz1Δ* mutant exhibited a dramatic increase in micafungin tolerance when pre-exposed to manogepix (Fig. 4). Checkerboard assays were utilized to determine whether manogepix increases resistance to micafungin in *crz1Δ* mutants. Manogepix did not antagonize micafungin in *crz1Δ* mutants (Fig. 5). However, manogepix did antagonize micafungin in wild-type parent strain, possibly due to calcineurin and Crz1-dependent expression of target genes such as *FKS2*. These findings suggest that calcineurin signaling is a key driver of tolerance and persistence in *C. glabrata*, with a portion of its effects mediated by Crz1 and another portion mediated by unknown effectors.

**Figure 4.**
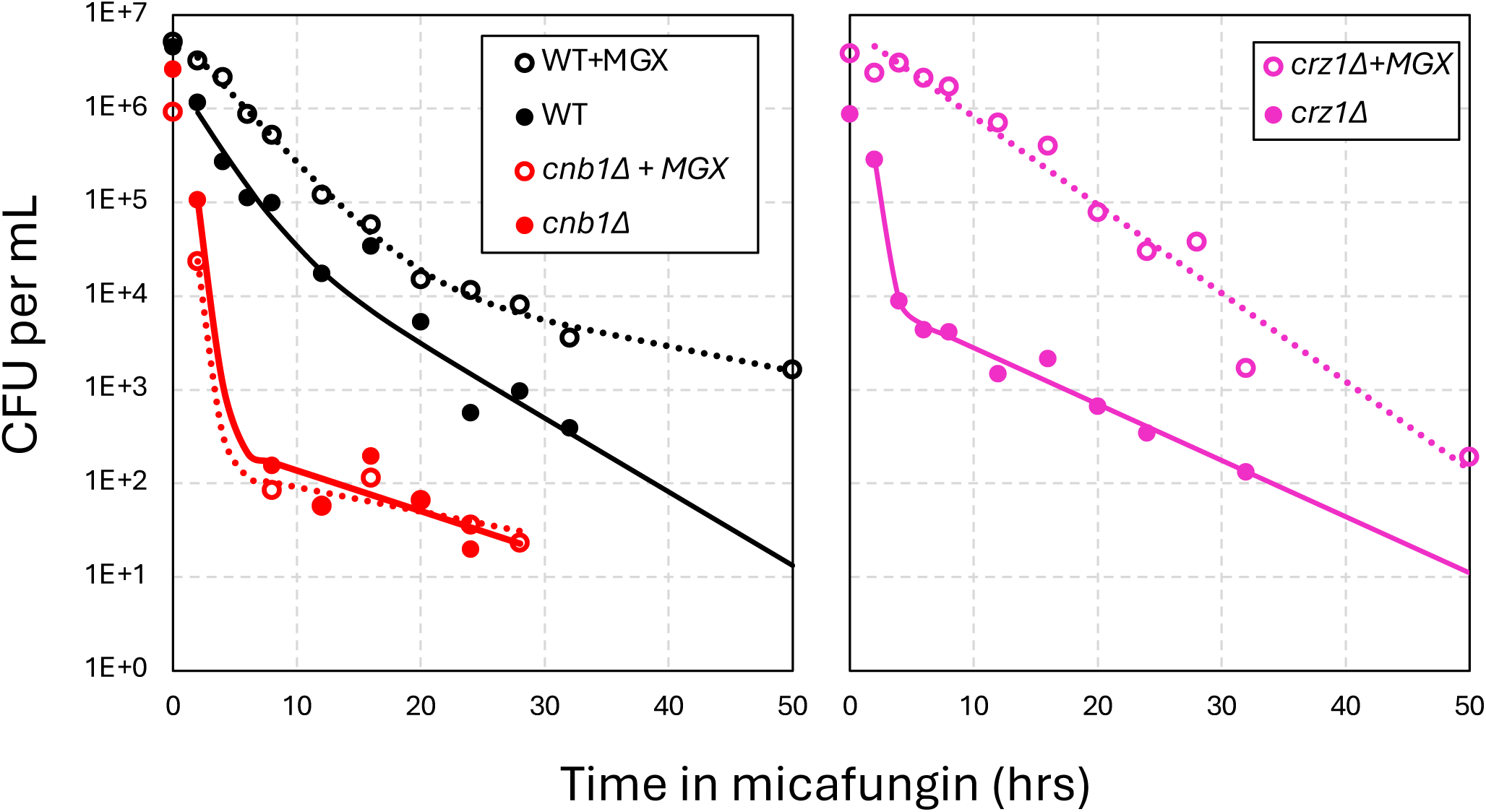
Manogepix activation of calcineurin induces micafungin tolerance and persistence independent of Crz1. Wild-type (black), *cnb1Δ* (red), and *crz1Δ* (purple) strains were grown to saturation and diluted 25-fold into fresh medium containing (dashed lines) or lacking (solid lines) manogepix (0.6 µg/mL). After shaking at 30°C for 2 hours, cultures were diluted 2-fold in fresh medium containing micafungin (125 ng/mL) and analyzed in time-kill experiments as described in Fig. 1. The averages of four biological replicates (symbols) were used for curve fitting (smooth curves).

**Figure 5.**
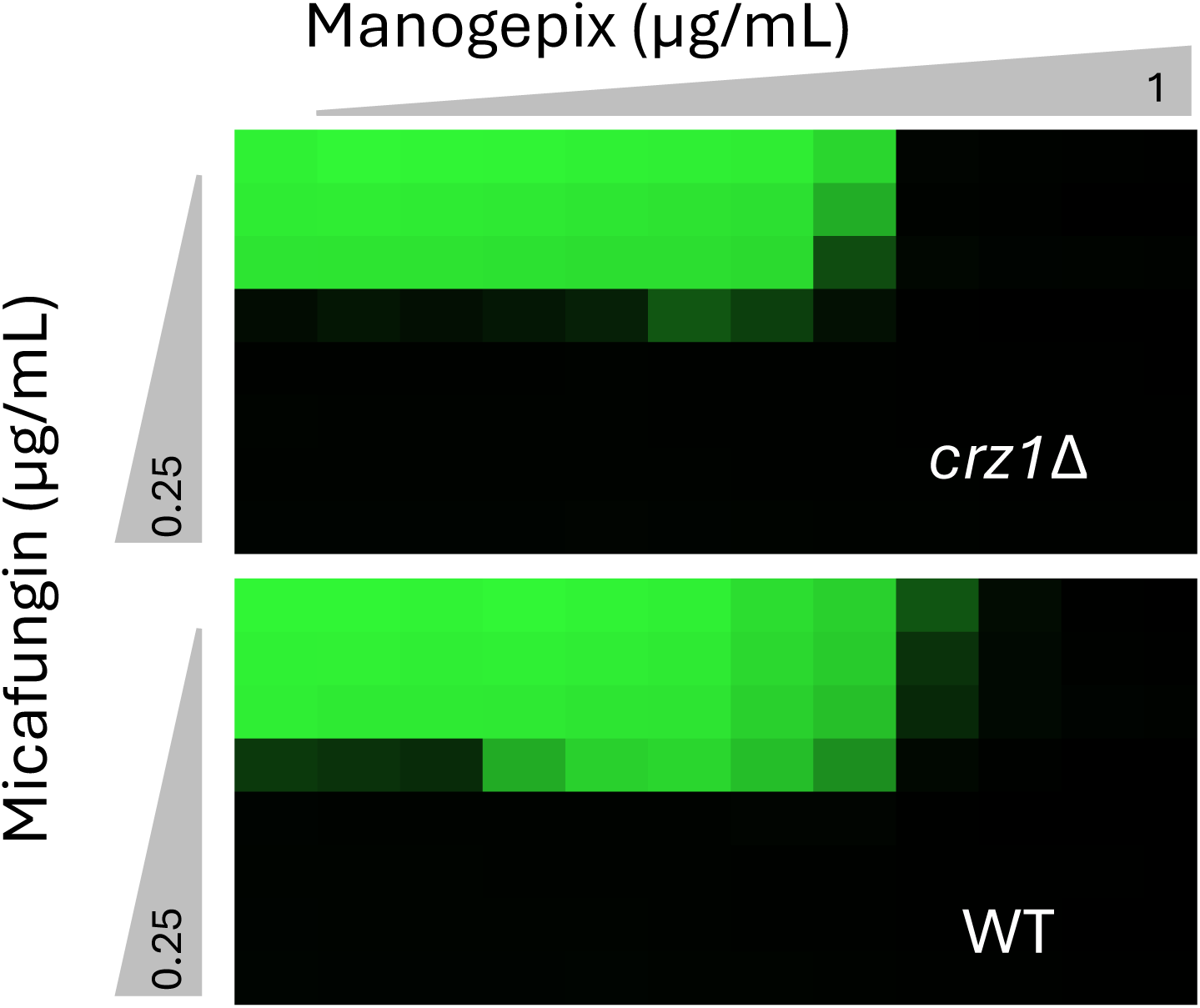
Mannogepix activation of calcineurin induces micafungin resistance through Crz1 and Fks2. Growth (green) of wild-type and *crz1Δ* strains was measured after incubation for 24 hr in medium containing varying concentrations of mannogepix and micafungin. Similar effects were seen in two additional replicates.

### Chronic Activation of Calcineurin increases tolerance and persistence

Some of the genes with positive tolerance/persistence Z-scores in the Tn-seq screen may generate chronic cellular stresses that pre-activate calcineurin when disrupted with transposons. One such gene is *FKS1* (Z = 3.38), which encodes the major catalytic subunit of glucan synthase (25, 39, 40). The *fks1Δ* mutants are FK506-sensitive because they depend on calcineurin signaling, Crz1, and elevated expression of *FKS2* for viability (25). When tested in time-kill assays, *fks1Δ* mutants exhibit a large increase in tolerance compared to BG14 and the parent strain (Fig. 6, Left). Thus, chronic pre-activation of calcineurin through Fks1-deficiency may increase tolerance and persistence much like the acute activator, manogepix.

**Figure 6.**
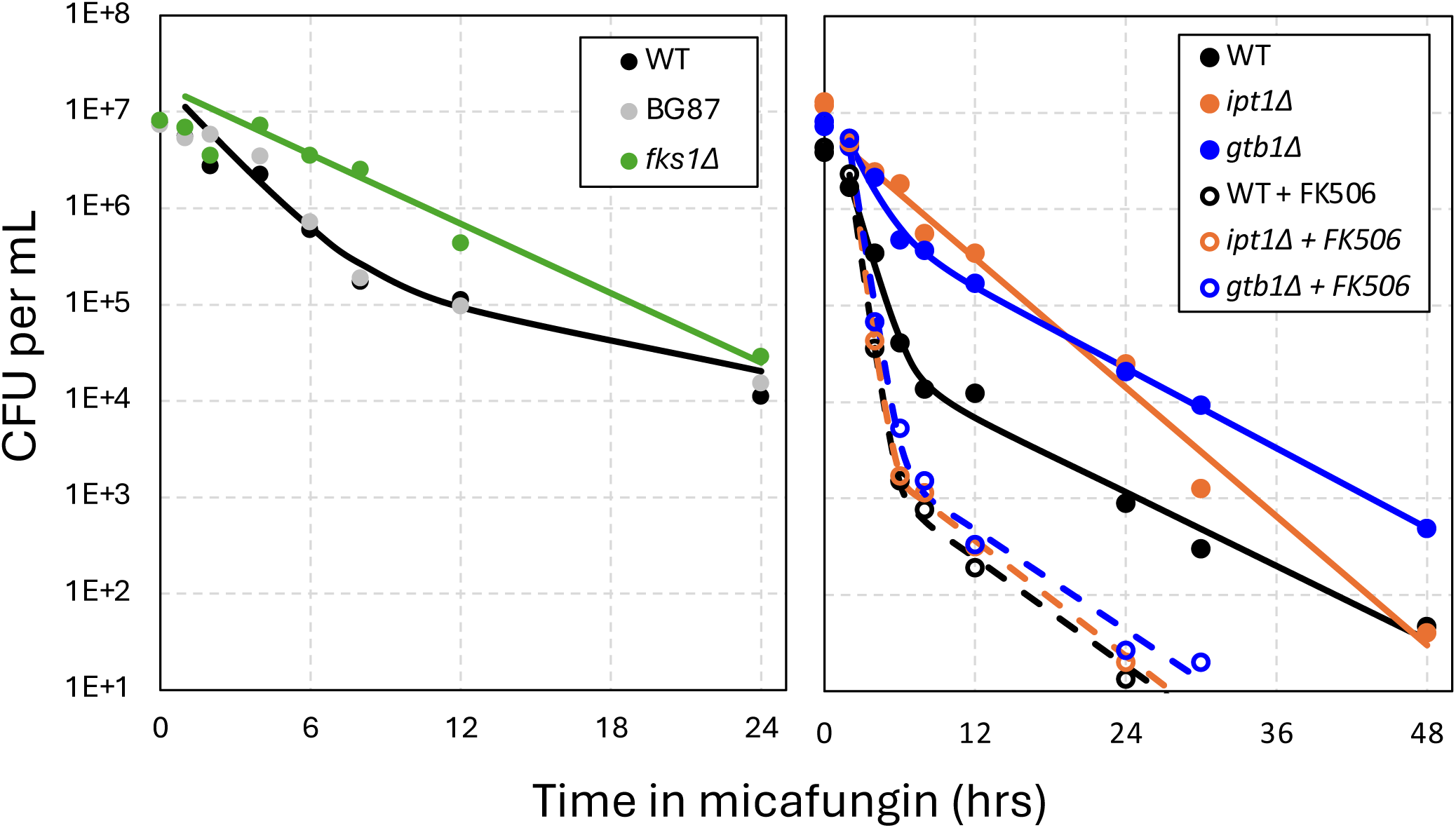
Genetic activation of calcineurin increases tolerance and persistence. The *fks1Δ* (green), *ipt1Δ* (orange), and *gtb1Δ* (blue) mutants were generated and tested in time-kill experiments as described in Fig. 1 in the absence (smooth curves) and presence (dashed curves) of FK506. Averages of four biological replicates were used to generate the curve fits.

Several genes encoding ER proteins (e.g. *CNE1*, *GTB1*, *KEG1*, *SKN1*, *VMS1*) exhibited strongly increased tolerance/persistence Z-scores (Table S1, Fig. 2). Knockout mutants of *GTB1* were generated and tested in time-kill experiments in the presence and absence of FK506. The *gtb1Δ* mutants exhibited elevated tolerance to micafungin that were strongly blocked by FK506 (Fig. 6, Right). The *IPT1* gene, which was also highly significant in the Tn-seq screen (Z = 13.0), encodes an enzyme in the Golgi complex that synthesizes the abundant sphingolipid M(IP2)C. In time-kill experiments, an *ipt1Δ* mutant exhibited elevated tolerance and persistence that was likewise blocked by FK506 (Fig. 6, Right). Heteroresistance to micafungin was not increased in the *fks1Δ* mutant (Fig. S6). These findings suggest that many negative regulators of calcineurin signaling may be among the list of genes with positive Z-scores in micafungin tolerance/persistence screens. However, other gene deficiencies may impact tolerance and persistence through calcineurin-independent effects.

### HOG signaling pathway negatively regulates tolerance and persistence independent of calcineurin

Calcineurin undoes the effects of serine/threonine protein kinases on shared substrates. Two genes encoding protein kinases (*FPK1*, *SSK2*) exhibited strongly positive Z-scores in the tolerance/persistence screen and therefore could produce phosphoproteins that are directly targeted by calcineurin. However, they were also strongly positive for micafungin resistance (Fig. 2). *SSK2* encodes a MAPKKK that phosphorylates and activates Pbs2, a MAPKK, that in turn phosphorylates Hog1, a MAPK, that is well known to respond to high-osmolarity signals (41). Though *PBS2* and *HOG1* were near zero in the Tn-seq screens, an upstream regulator of *SSK2* (*SSK1*) also exhibited positive tolerance/persistence and resistance Z-scores. Knockout mutants lacking the *SSK2*, *PBS2*, and *HOG1* genes in the BG14 background were analyzed in time-kill experiments using 8-fold higher doses of micafungin to compensate for their mild resistance to echinocandins. All three mutants exhibited elevated tolerance relative to the wild-type control in the presence of FK506 (Fig. S7, Fig. 7). These results indicate the HOG signaling pathway normally negatively regulates tolerance and persistence independent of calcineurin. In the absence of FK506, calcineurin signaling strongly increased tolerance and persistence in all three mutants (Fig. 7, Fig. S7), which indicates that calcineurin promotes tolerance and persistence independent of HOG signaling. The effects of *hog1Δ* mutations were also observed in a *crz1Δ* mutant background (Fig. 7). Thus, HOG signaling can negatively regulate tolerance and persistence independent of Crz1 and calcineurin.

**Figure 7.**
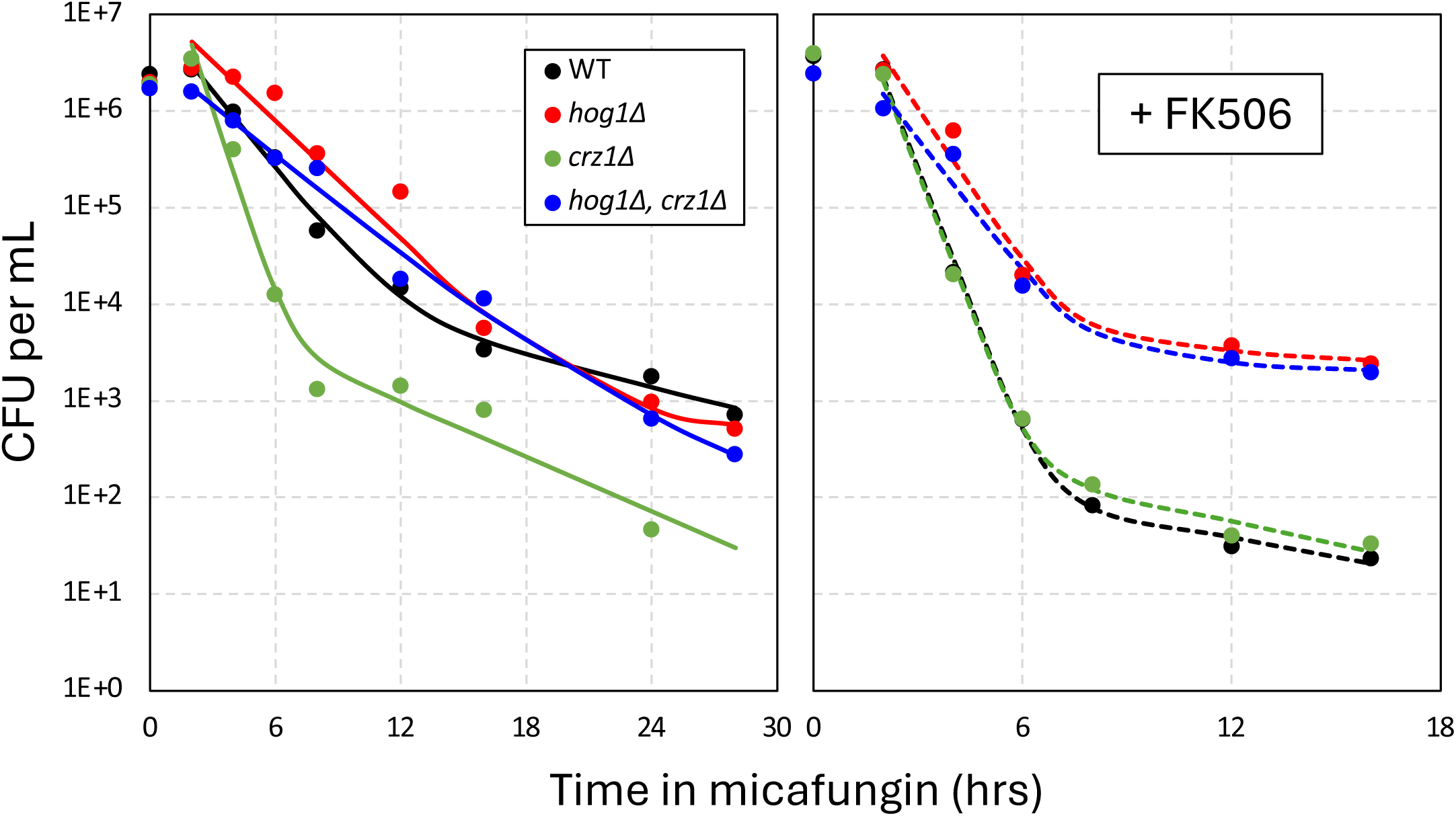
Hog1 diminishes tolerance and persistence independent of calcineurin and Crz1. The *hog1Δ crz1Δ* double knockout mutant (blue), the single knockout mutants (green, red), and wild-type control strain (black) were analyzed in time-kill experiments as described in Fig. 1 with the effects of FK506 illustrated separately (right panel). Averages of four biological replicates were used to generate the curve fits.

### The CBS138 strain of *C. glabrata* utilizes calcineurin but not HOG signaling to regulate micafungin tolerance and persistence

The CBS138 strain of *C. glabrata* naturally carries a mutation that inactivates the *SSK2* component of the HOG signaling pathway (41). This mutation may be adaptive by providing enhanced tolerance and persistence, resulting in infections that are more difficult to treat. To begin investigating this possibility, time-kill experiments were performed on *pbs2Δ*, *cna1Δ, cnb1Δ, and crz1Δ* mutants in the CBS138 strain background. Unlike its effects in the BG14 strain background, the *pbs2Δ* mutation had no significant impact on tolerance and persistence in the CBS138 strain background, which exhibited somewhat elevated tolerance and persistence even in the presence of FK506 (Fig. S8). The effects of *cna1Δ* and *cnb1Δ*, were conserved in CBS138, however the effects of *crz1Δ* were small relative to BG14 (Fig. S9). The natural deficiency of HOG signaling and other polymorphisms in CBS138 may contribute to its enhanced antifungal tolerance. Interestingly, a recent survey of CBS138 and dozens of additional *C. glabrata* strains revealed considerable variation in the number of persister-like cells observed after micafungin exposure (30).

### Calcineurin signaling promotes tolerance and persistence to micafungin, but not amphotericin B, in *C. albicans*

To test whether calcineurin drives tolerance and persistence in *C. albicans*, micafungin time-kill experiments were performed on wild-type strain SC5314 in the presence and absence of FK506. Biphasic kinetics of cell death were observed in both scenarios while the loss of calcineurin signaling decreased the half-life of susceptible cells approximately 1.4-fold and decreased the number of persister-like cells approximately 100-fold (Fig. 8, black symbols). Similar results were obtained previously using *cna1Δ/Δ* mutants of *C. albicans* instead of FK506 (42). Curiously, the half-life of persister-like cells in *C. albicans* was much longer than that of *C. glabrata*, indicating that the two species may regulate the process in different ways.

**Figure 8.**
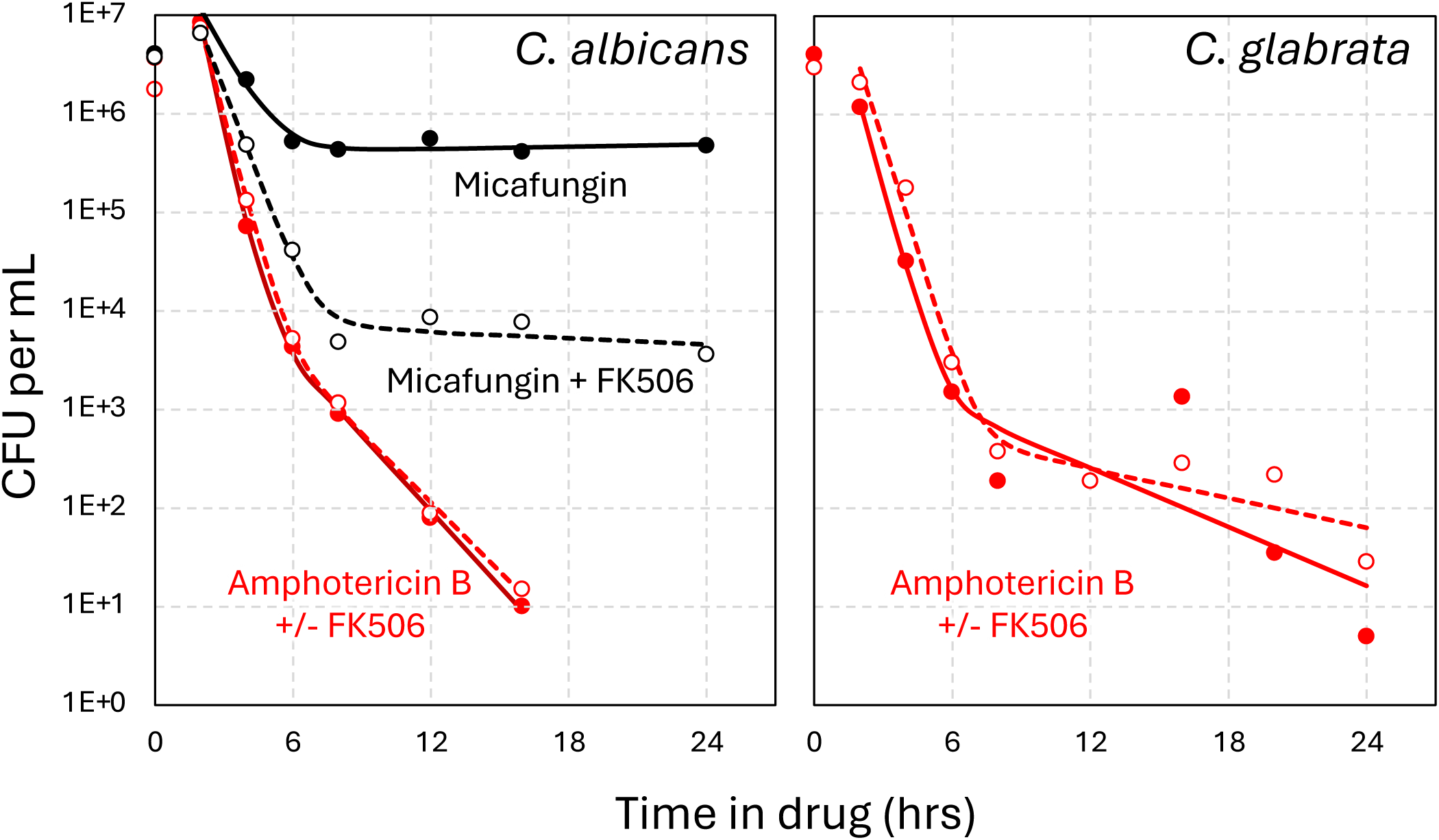
Calcineurin induces micafungin tolerance and persistence in *C. albicans* but has no impact on amphotericin B. Wild-type strains of *C. albicans* (left) and *C. glabrata* (right) were analyzed by time-kill experiments using 125 ng/mL micafungin (black lines) or 10 µg/mL amphotericin B (red lines) in the absence or presence of FK506 (dashed lines) as described in Fig. 1. Averages of four biological replicates were used to generate the curve fits.

Amphotericin B, which kills fungal cells by depleting ergosterol and creating pores in the plasma membrane (43), was also found to kill *C. albicans* and *C. glabrata* with biphasic kinetics after a brief lag (Fig. 8, red symbols). The half-life of the susceptible population was 2 to 2.5-fold shorter than observed with micafungin and, in both species, the killing kinetics were not altered by FK506 (dashed lines). These findings suggest that calcineurin specifically drives tolerance to echinocandins, but not amphotericin B in these species and conditions.

## DISCUSSION

Though there is concern that tolerance and persistence will contribute to therapeutic failure, relapse, and perhaps even the acquisition of resistance (30, 44), the regulatory mechanisms behind these phenomena have been understudied in fungal pathogens of humans. This study shows that calcineurin signaling is a major driver of tolerance and persistence to echinocandins in *C. glabrata* and *C. albicans*, in addition to driving resistance. Calcineurin and Crz1 produce mild resistance to echinocandins in *C. glabrata* by increasing expression of the drug target Fks2 (25). When strong resistance mutations in *FKS2* arise in patients, calcineurin inhibitors greatly diminish their impact *in vitro* (25, 39). Here we show that calcineurin signaling also increases tolerance and persistence largely independent of *CRZ1, FKS2*, and *RCN2*. Though the target of calcineurin that promotes tolerance and persistence has not yet been identified, the HOG signaling pathway was not required. FK506 still lowered tolerance and persistence in *ssk2Δ*, *pbs2Δ*, *hog1Δ*, and *crz1Δ hog1Δ* mutants. However, all these mutants still exhibited elevated tolerance and persistence in the absence of calcineurin signaling, again through unknown effectors (Fig. 7, Fig. S7). Interestingly, the HOG signaling pathway is naturally polymorphic in different strains of *C. glabrata*, contributing to variation in the resistance to micafungin (21) and other stressors (41), as well as variation in tolerance and persistence within the species. Multiple strains of *C. glabrata* may be needed to obtain a complete picture of resistance, tolerance, and persistence mechanisms in this diverse species. As calcineurin also increases echinocandin tolerance in *C. albicans* and *A. fumigatus* (45, 46), the mechanism may be broadly conserved among diverse fungal pathogens.

A recent study concluded that mitochondrial dysfunction regulates echinocandin tolerance in *C. glabrata* (47). However, that study did not make use of time-kill experiments and extensively utilized low-resolution dose-kill experiments, where CFU are quantified at a single time point (24 hr) after exposure to varying doses of echinocandins (23). The study also used “mitochondrial inhibitors” that either have multiple cellular targets (sodium azide) or have no targets at all in *C. glabrata* (diphenyleneiodonium chloride, rotenone) and did not match several mutants lacking mitochondrial functions. Nevertheless, after 24 hr exposure to high doses of caspofungin, several mitochondrial mutants produced wild-type CFU, consistent with our mitoribosome-deficient *mrp20Δ* mutant, which exhibited little or no changes in tolerance and persistence (Fig. S4). Additionally, hundreds of mitochondrial gene deficiencies caused by transposon insertions exhibited near-zero tolerance/persistence Z-scores and strongly positive resistance Z-scores (Fig. 2) due, in part, to their activation of Pdr1 signaling and elevated expression of lipid flippases that weakly increase resistance to most echinocandins (21). This study highlights the benefits of high resolution time-kill experiments are performed at supra-MIC doses and coupled with conventional mathematical modeling.

A noteworthy observation of this study is that the estimated number of persister-like cells (i.e. persistence) correlated with the half-life of susceptible cells (i.e. tolerance) in all the mutant strains tested here. By increasing tolerance in the majority of cells in the population, calcineurin signaling may buy time for persister-like cells to form *de novo* through other regulatory processes. In the absence of calcineurin signaling, tolerance was low and persister-like cells were rare, but still readily detectable. When calcineurin was pre-activated by mutations or inhibitors that target the ER (e.g. manogepix), tolerance and persistence were elevated (Fig. 4). The ability of FK506 to block these effects suggests that ongoing calcineurin signaling is necessary to maintain tolerance during micafungin exposure and enable persistence to arise. In addition to increasing micafungin tolerance and persistence independent of Crz1, manogepix increased micafungin resistance through Crz1-dependent processes (Fig. 5). Manogepix, a fungistat that is currently in phase 2 clinical trials to treat a range of fungal infections (38), may therefore produce adverse drug-drug interactions with echinocandins. However, when combined with FK506, manogepix becomes fungicidal against *C. glabrata* and a wide range of fungal pathogens (36, 48). The role of calcineurin in manogepix tolerance may overlap with previously studied calcineurin-dependent tolerance to tunicamycin and to azole-class antifungals, which target N-glycosylation and ergosterol biosynthesis in the ER, respectively (49).

Other environmental conditions may effect calcineurin and HOG signaling pathways leading to impacts on echinocandin tolerance and persistence. Engulfment of *C. glabrata* cells into phagosomes of macrophages produced elevated micafungin tolerance and persistence (31). Tolerance to amphotericin B was also observed for engulfed *C. glabrata* cells (31). It will be interesting to determine whether calcineurin and/or HOG signaling govern these processes, or whether the longer lifespans of engulfed cells arise simply through slower growth rates or perhaps slower drug access to this intracellular compartment. Calcineurin signaling promotes virulence of *C. glabrata* (27, 28)*, C. albicans* (42, 50, 51), and other fungal pathogens (45, 52–54) in mouse models of systemic candidiasis. In *C. glabrata*, the effects of calcineurin were independent of Crz1 (27, 28). A new interpretation of those findings is that host environments produce toxins or hostile conditions to the pathogens and generate stresses, which might normally activate calcineurin, induce tolerance and persistence behaviors, and promote fungal cell survival during infection. Manogepix treatment may further activate calcineurin and augment tolerance and persistence mechanisms, while still exerting a fungistatic effect. On the other hand, FK506 treatment may lower the defenses of *C. glabrata* and enable much more rapid and complete killing by host-derived assaults as well as echinocandins. Non-immunosuppressive analogs of FK506 that specifically target fungal calcineurin could be highly effective antifungals alone or in combination with existing antifungals (55). Though the immunosuppressive effects of FK506 would preclude long-term use of this compound as an antifungal co-drug, our findings raise the possibility that short-term or bolus co-administration of FK506 during standard echinocandin therapies could improve outcomes without producing sustained immunosuppression.

In a mouse model of invasive candidiasis by *C. albicans*, the efficacy of fluconazole was increased by co-administration of FK506 (56). Tolerance to fluconazole and other inhibitors of lanosterol desaturase (Erg11) in *C. glabrata*, *C. albicans*, and other yeast species has long been known to depend on calcineurin signaling, but not Crz1 (27, 28, 49, 52). Calcineurin also promoted tolerance to terbinafine in *C. albicans* (57), tunicamycin and dithiothreitol in *S. cerevisiae* (58, 59), and to SDZ 90-215 in all these species (60). These compounds all block essential enzymes in the ER or Golgi complex of the fungal cells and are considered fungistatic in wild-type cells and fungicidal in calcineurin-deficient cells. Because the wild-type yeasts do not lose viability in the presence of these compounds, and often continue to replicate slowly for a few doublings, the degree of tolerance conferred by calcineurin signaling is difficult to quantify with the mathematical models employed in this study. The effectors of calcineurin responsible for tolerance to these fungistats remain unknown. Therefore, it is not yet possible to determine whether the same effectors govern tolerance to these fungistats as well as the fungicidal echinocandins. Our Tn-seq screen has produced dozens of candidate genes that could mediate the effects of calcineurin and potentially serve as targets for development of non-immunosuppressive strategies to increase the efficacies of our most important antifungals.

Though calcineurin promoted tolerance to echinocandins and azoles, it did not seem to alter the biphasic kinetics of survival during exposure to amphotericin B in *C. glabrata* or *C. albicans* or *S. cerevisiae* (61). Amphotericin B kills growing and non-growing fungal cells by sponging essential ergosterols from the plasma membrane and generating lethal pores (43). The procedures outlined in this study could be adapted for studies of amphotericin B tolerance and persistence mechanisms. As new derivatives of amphotericin B with dramatically less animal toxicity are being developed (62), a more complete understanding of polyene resistance, tolerance, and persistence mechanisms in fungi could further increase their therapeutic index.

## METHODS

### *Candida* strains and plasmids

All strains and their sources are listed in Table S2. *C. glabrata* strains were derived from BG14, a *ura3Δ* derivative of wild-type strain BG2 (63). Individual gene knockouts were constructed using the PRODIGE method (64) where coding sequences were replaced with the coding sequence of *URA3* from *S. cerevisiae* plasmid pRS406 (65) using oligonucleotides listed in Table S3. Colony PCR was used for screening validation. The *RCN2* gene was PCR amplified from genomic DNA of strain BG14 and cloned into the centromeric plasmid pCN-PDC1 (66) to generate pCN-PDC1-RCN2 and authenticated via Sanger sequencing.

### Antifungal Drugs

Micafungin (Cat. #18009) and Amphoteracin B (Cat. #11636) were obtained from Cayman Chemicals, caspofungin (Cat. #S3073), tactrolimus (FK506) (Cat. #S5003), and mannogepix (E1210) (Cat. #S0491) from SelleckChem, and ibrexafungerp from Scynexis.

### TN-seq screen for genes that regulate micafungin tolerance and persistence

Pool-3 of *Hermes-NAT1* insertion mutants in strain BG14 that had been studied previously for resistance to echinocandins (21) was thawed from storage at −80°C, grown to saturation, then diluted into fresh SCD-0 medium containing 64 µg/mL micafungin. After 0 and 6 hr of shaking at 30°C, the culture was sampled, chilled on ice, washed once with chilled SCD-0 medium, resuspended in SCD-0, and shaken at 30°C for another 48 hr to allow regrowth of surviving cells. The cells were pelleted, washed, and then genomic DNA was extracted from using the Quick-DNA Fungal/Bacterial Miniprep Kit from Zymo Research (Cat. #D6005). gDNA was fragmented by sonication, repaired, ligated to adapters, size-selected, and then insertion sites were PCR amplified and sequenced on an Illumina MiSeq instrument as described previously (21). FastQ files were demultiplexed using CutAdapt (67) and aligned to the BG2v1 reference genome (68) using Bowtie2 (69). Mapped reads with low quality (Q<20) or mismatches at position 1 were discarded and the remainder were tabulated for each site in the genome. The total number of sequence reads in each annotated gene were tabulated and Z-scores were calculated for both time points with respect to the starting pool and with respect to each other as before (21). Gene Ontology analysis was performed on subsets of analyzed genes using GOrilla in *S. cerevisiae* mode (33).

### Time-kill assays and analyses

At least four single colonies of each *C. glabrata* strain were picked and grown to saturation for 72 hours in SCD-0 media at 30°C. Microscopic observations indicated that the *C. glabrata* cells were monodispersed and not detectably clumped or aggregated. Each replicate culture was diluted 1:50 into fresh media containing antifungals, and shaken at 30°C. Aliquots were periodically removed, serially diluted in SCD-0 medium, and immediately plated onto SCD-0 agar medium using a sterile pinning device or by pipetting. Plates were incubated at 30°C until individual colonies were visible under a dissecting microscope at 10x magnification. Visible colonies were counted manually at one dilution for each sample and CFU per mL of undiluted culture was calculated. Replicates were averaged arithmetically and charted. For curve-fitting, the CFU were log transformed and fit to the equation y = log(m1*exp(-m2*x) + m3*exp(-m4*x)) by non-linear regression in Kaleidagraph (v5.04, Synergy) after exclusion of data points within the lag phase. Parameters m1 and m3 represent sizes of the non-persister and persister cell populations, respectively. Parameters m2 and m4 represent kill constants that were converted to half-lives (= ln2/m2, ln2/m4) of the two populations. Smooth curves were generated using these parameters and overlayed onto the CFU data points. The same procedure was applied for analysis of *C. albicans* except that YPD medium was used.

### Population Analysis Profiling (PAP) Assay

Single colonies were grown to saturation for 72 hours in SCD-0 media at 30°C. The saturated cultures were diluted 1:50 into fresh SCD-0 media and then serially diluted 1:5. 2 µL of each dilution was spotted onto agar YPD media containing varying doses of micafungin (15 ng/mL to 0.234 ng/mL). Alternatively, when viability was very low, 200 µL of undiluted cultures were plated. Plates were incubated for 24 hours at 30°C and single colonies were counted manually. CFU per mL original culture was then calculated and replicates were averaged.

### Checkerboard Assay

Varying concentrations of micafungin and mannogepix were prepared in SCD-0 and aliquoted to their respective columns and rows in a 96-well dish. Three biological replicates of BG14 and *crz1Δ* strain were growth to saturation and diluted 3:2000 in fresh SCD-0 media before being aliquoted into 96-well plates one-third of the final well-volume for 1:2000 cell dilution. The 96-well plates were incubated for 24 hours at 30°C, mixed, and optical density at 600 nm was measured (Accuris SmartReader 96T).

## Data Availability

The authors affirm that all data necessary for confirming the conclusions of the article are present within the article, figures, tables, and repository. Raw sequencing reads used in this study were deposited at the NCBI Sequence Read Archive (SRA) with the BioProject ID PRJNA1304976. Micafungin resistance data (21) were obtained from SRA BioProject ID PRJNA1247003. A tabulation of the chromosomal coordinates and frequency of each mapped transposon insertion site (site count files) and a tabulation of the number of mapped transposon sites that fall within annotated gene boundaries (gene count files) are available upon request.

## ACKNOWLEDGMENTS

The authors thank Dr. Winston Timp for providing access to DNA sequencing instruments and Drs. Joseph Heitman, Karl Kuchler, and Alejandro de las Peñas for providing *C. glabrata* and *C. albicans* strains, and Dr. Winston Timp for providing access to critical instruments. This research was supported by grants from the National Institutes of Health (T32-GM007231 to the Cell, Molecular, Developmental Biology, and Biophysics doctoral, training program; R01-AI153414 to KWC).

---

**Fig S1.**
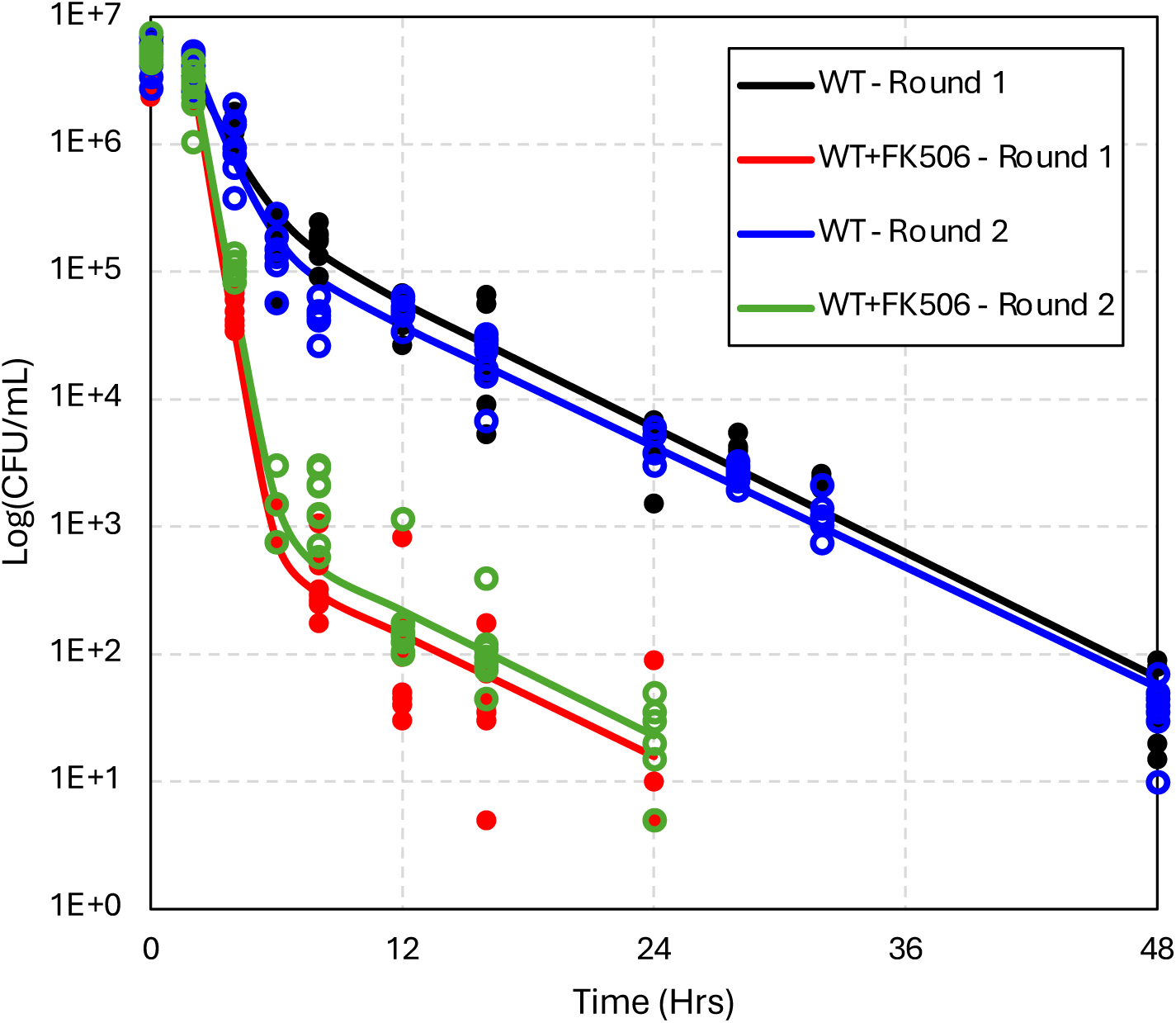
Biological replicates experience similar responses to micafungin and FK506 treatment. Eight single colonies of wild-type BG14 cells were grown to saturation for 72 hours, then diluted 50-fold into media containing micafungin (0.125 µg/mL) medium containing or lacking FK506 (1 µg/mL) and sampled as described in Fig. 1. Here, individual replicate cultures were counted and plotted (symbols) were plotted with curve fits imported from Fig 1.

**Fig S2.**
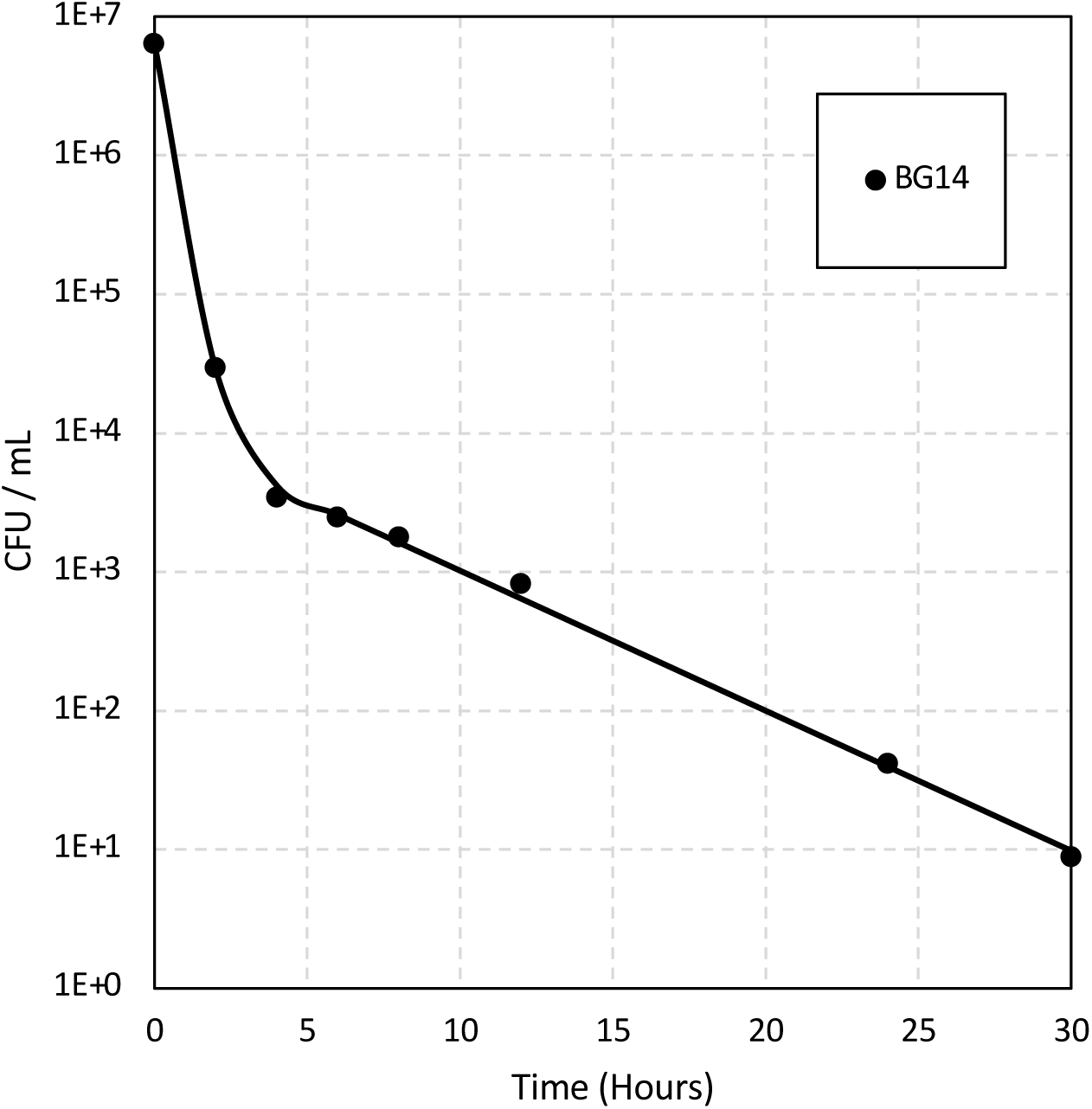
Log-phase populations exhibit no lag phase in time-kill experiments. Four single colonies were grown in fresh SCD medium for 24 hours until log phase phase was achieved. Time-kill assays with micafungin were performed as described in Fig 1.

**Figure S3.**
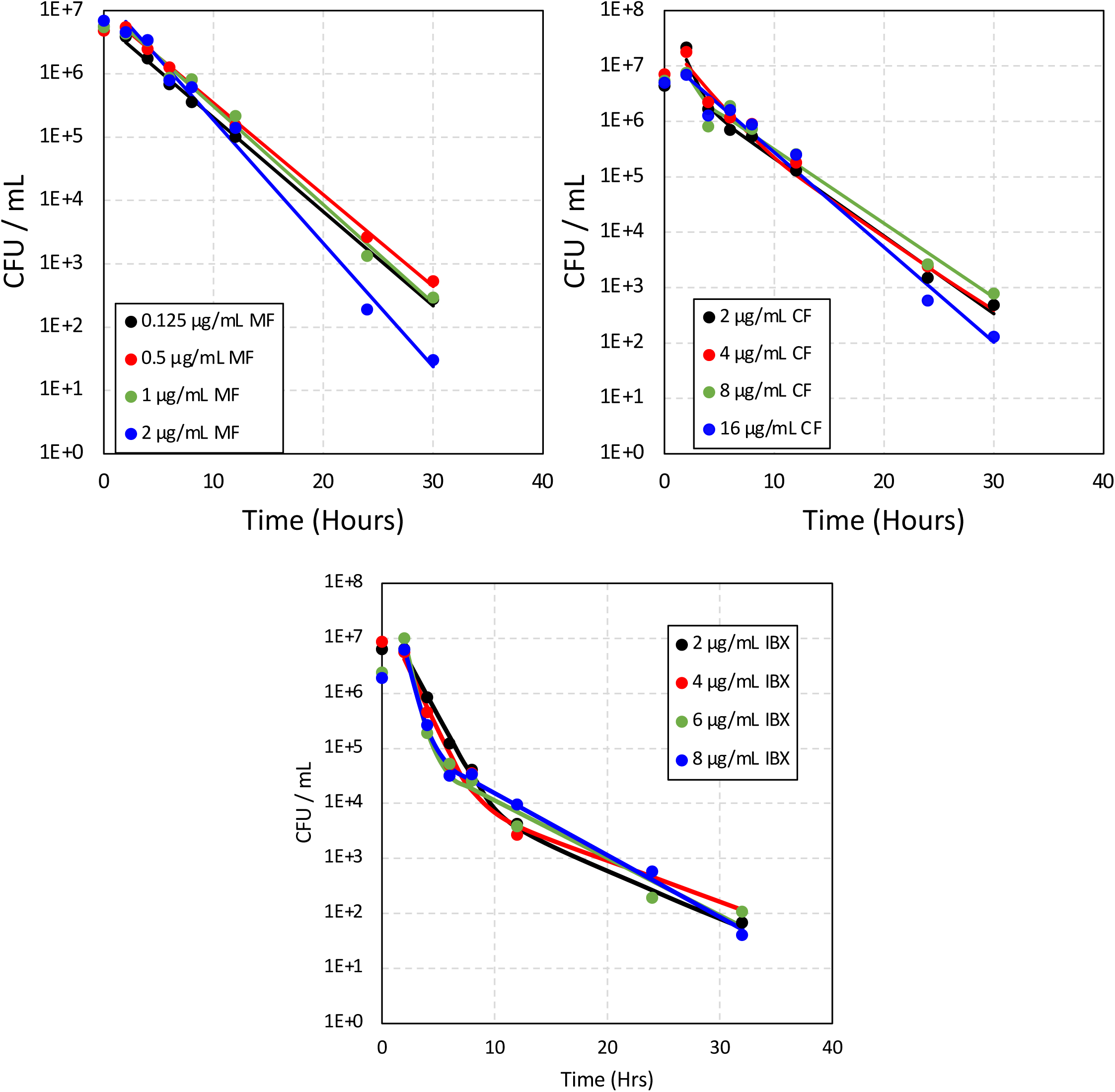
Tolerance and persistence to β-1,3-glucan synthase inhibitors is dose- and drug-independent. Wild-type cells were grown and treated with varying concentrations of micafungin (MF), caspofungin (CF), or ibrexafungerp (IBX) in as described Fig 1. Data points are the averages of four biological replicates and were used to generate the curve fits.

**Figure S4.**
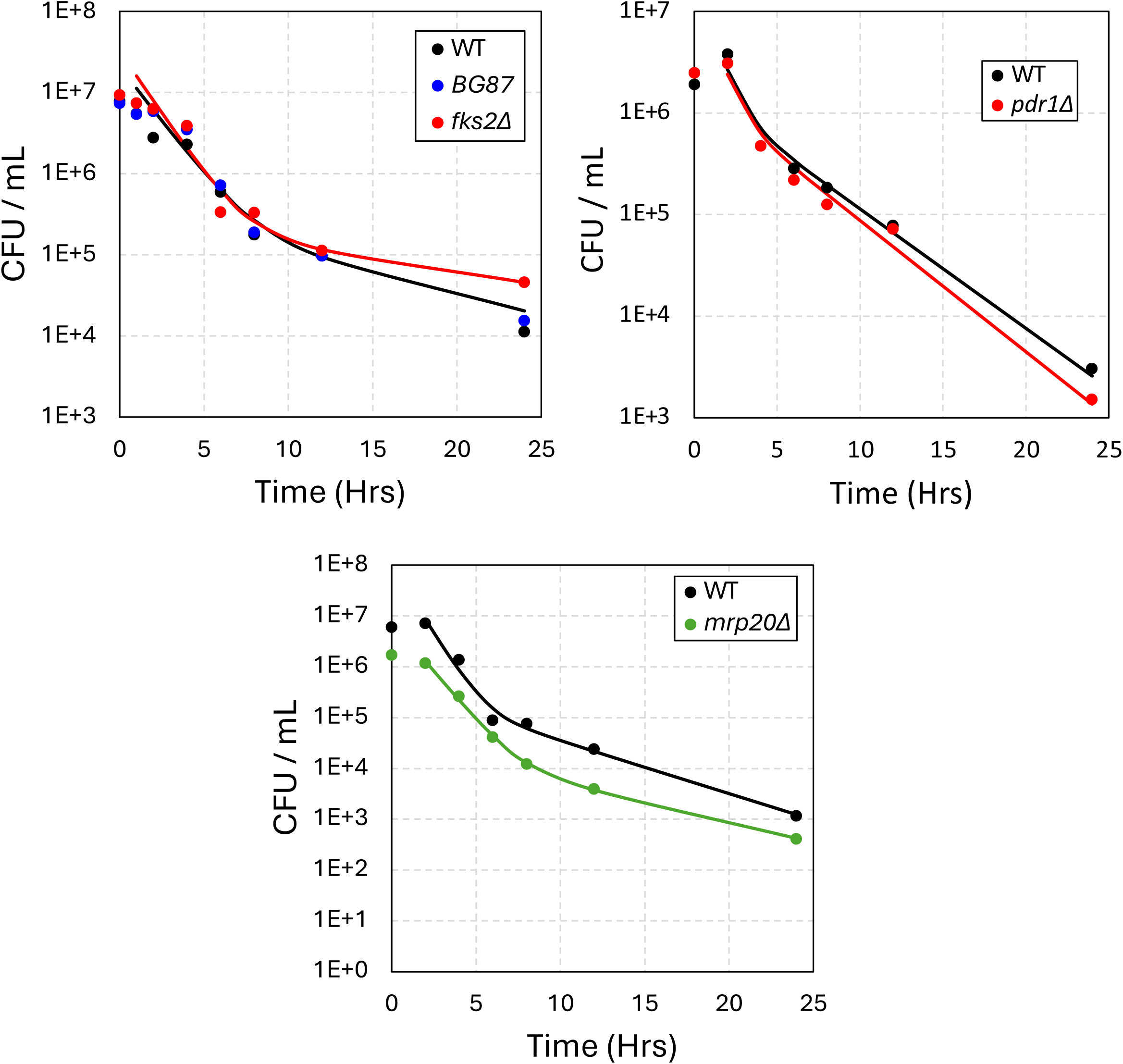
Strains associated with increased and decreased micafungin resistance do not alter tolerance or persistence. (Top Row) Strains were grown and treated with micafungin as described Fig 1. (Bottom) *mrp20Δ* mutant was treated with an 8X dose (1 µg/mL) of micafungin to compensate for resistance. Cultures were sampled and CFUs were calculated as described in Fig. 1. Data points are the averages of four biological replicates and were used to generate the curve fits.

**Figure S5.**
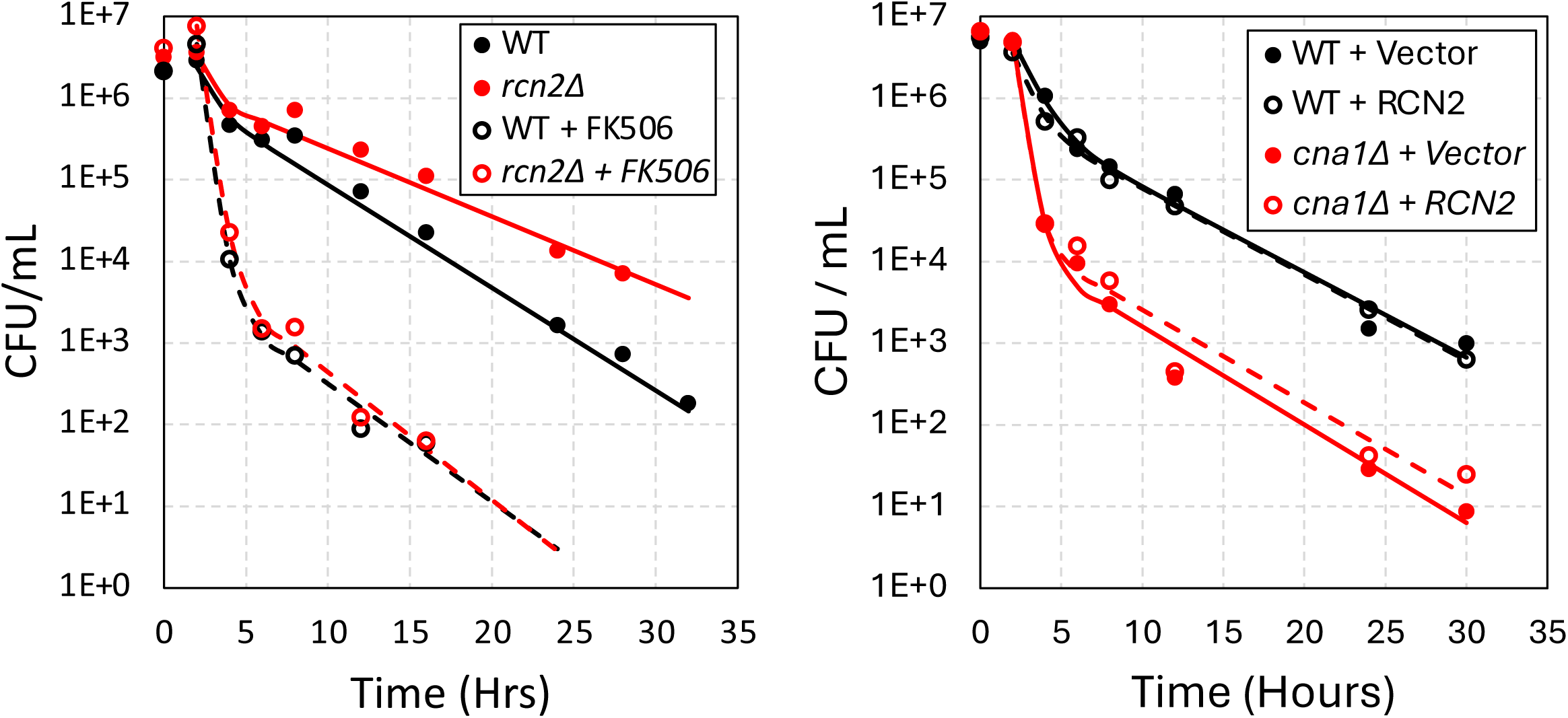
*RCN2* negatively regulates calcineurin in time-kill experiments. (Left) Wild-type and *rcn2Δ* strains were grown and treated with micafungin containing or lacking FK506 and time-kill experiments were performed as described Fig 1. Data points are the averages of four biological replicates and were used to generate the curve fits. (Right) Wild-type and *cna1Δ* strains were transformed with either empty plasmid (+ vector) or pCN-PDC1-RCN2 *(+ RCN2*). Single colonies were grown to saturation in SCD+NAT media for 72 hours to select for plasmid, then washed and diluted into fresh medium containing micafungin and and analyzed as described in Fig. 1. Data points are the averages of four biological replicates and were used to generate the curve fits.

**Figure S6.**
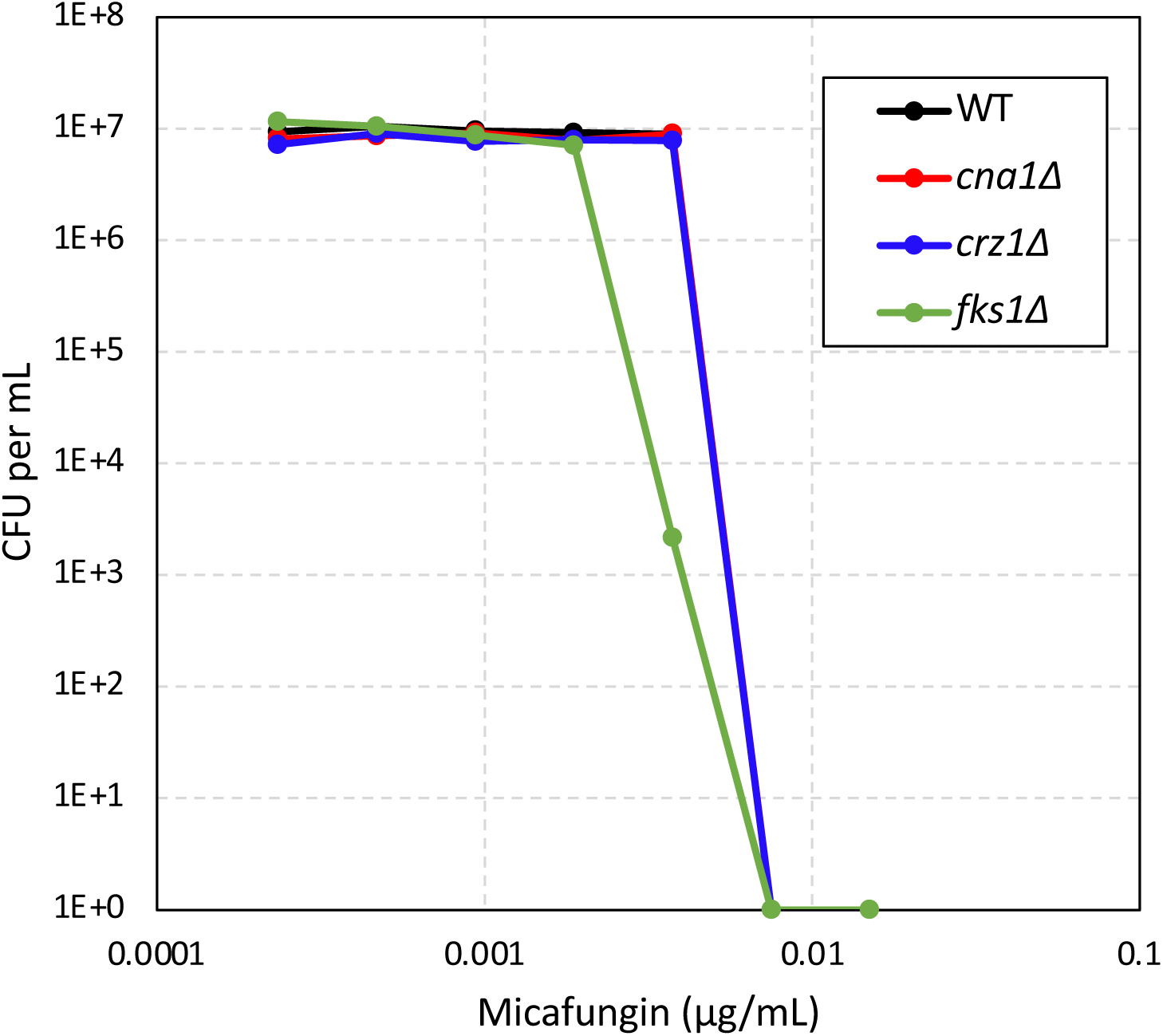
Heteroresistance to micafungin was not observed in *C. glabrata* strains. Wild-type (BG14) and mutant strains were grown to saturation for 72 hours, diluted into fresh SCD medium, and then plated on YPD agar media containing varying doses of micafungin (0 to 0.015 µg/mL). Plates were incubated for 24 hours at 30°C and colonies were counted using a dissecting microscope.

**Figure S7.**
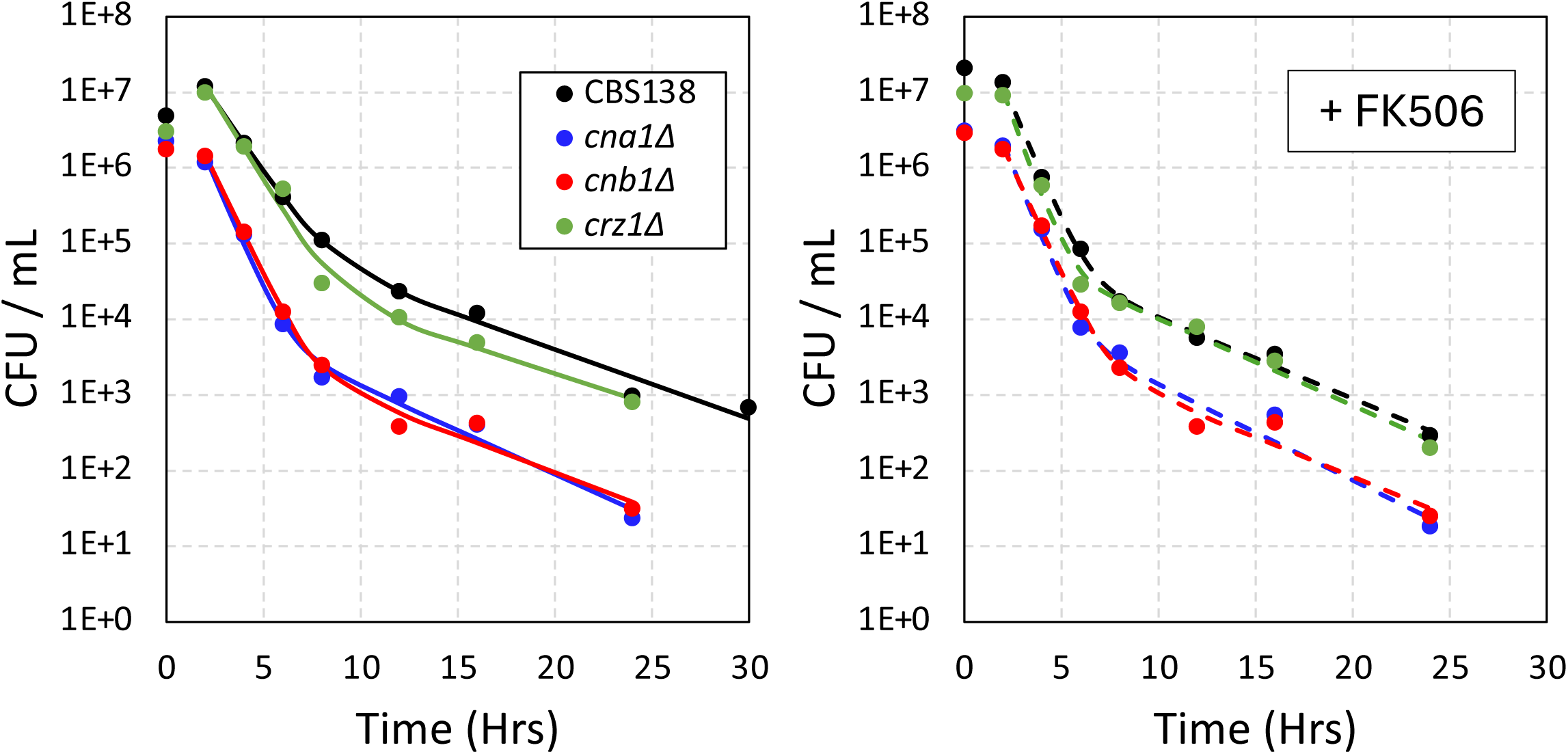
CN-dependent tolerance and persistence is conserved in the CBS138 strain of *C. glabrata*. Strains were grown and treated with micafungin containing or lacking FK506 as described Fig 1. Data points are the averages of four biological replicates and were used to generate the curve fits.

**Figure S8.**
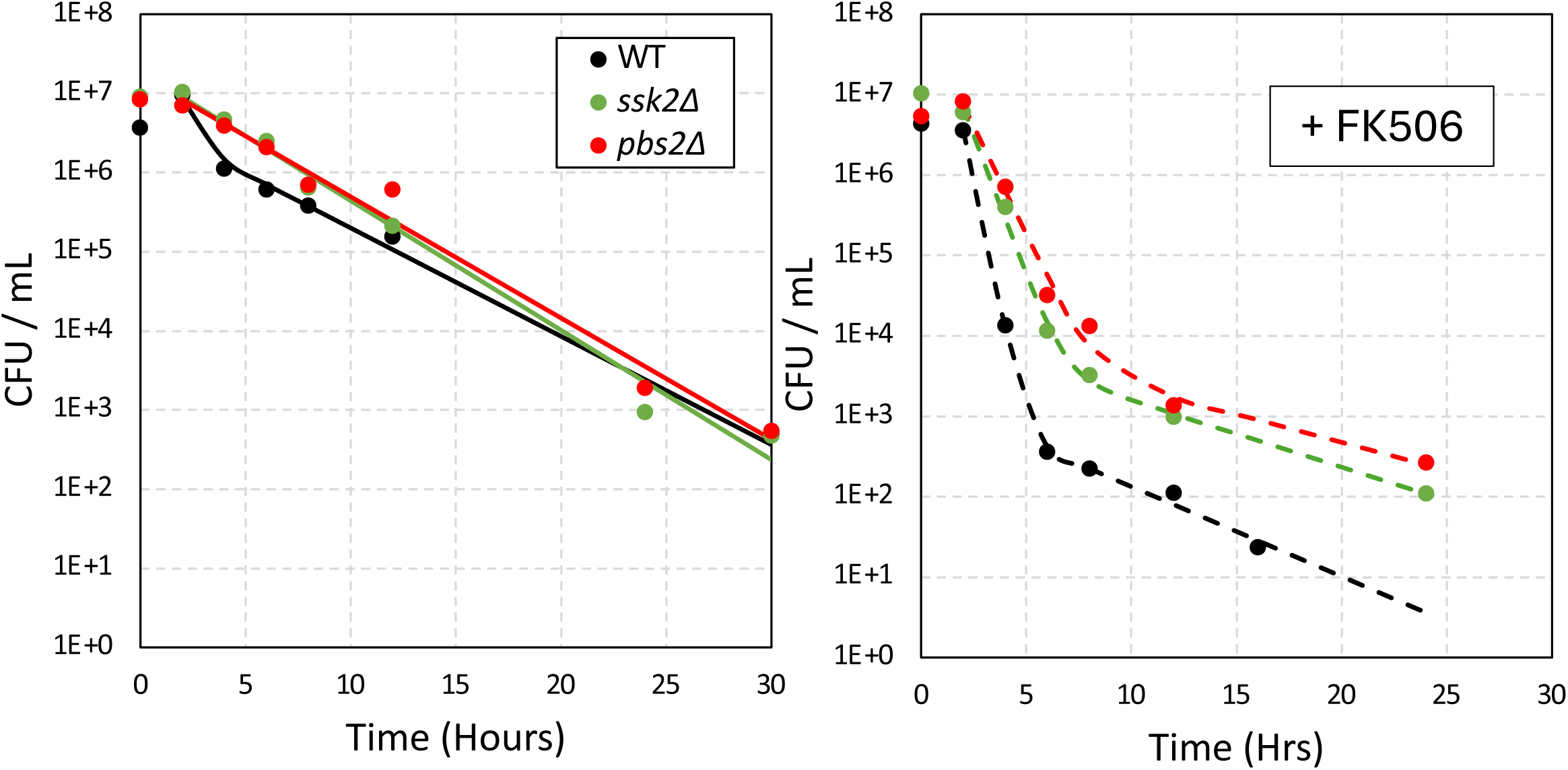
The HOG pathway negatively regulates tolerance and persistence independent of calcineurin. Strains were grown as describe in Fig. 1, treated with a higher dose of micafungin (1 µg/mL) containing or lacking FK506 (1 µg/mL), and sampled as previously described. Data points are the averages of four biological replicates and were used to generate the curve fits.

**Figure S9.**
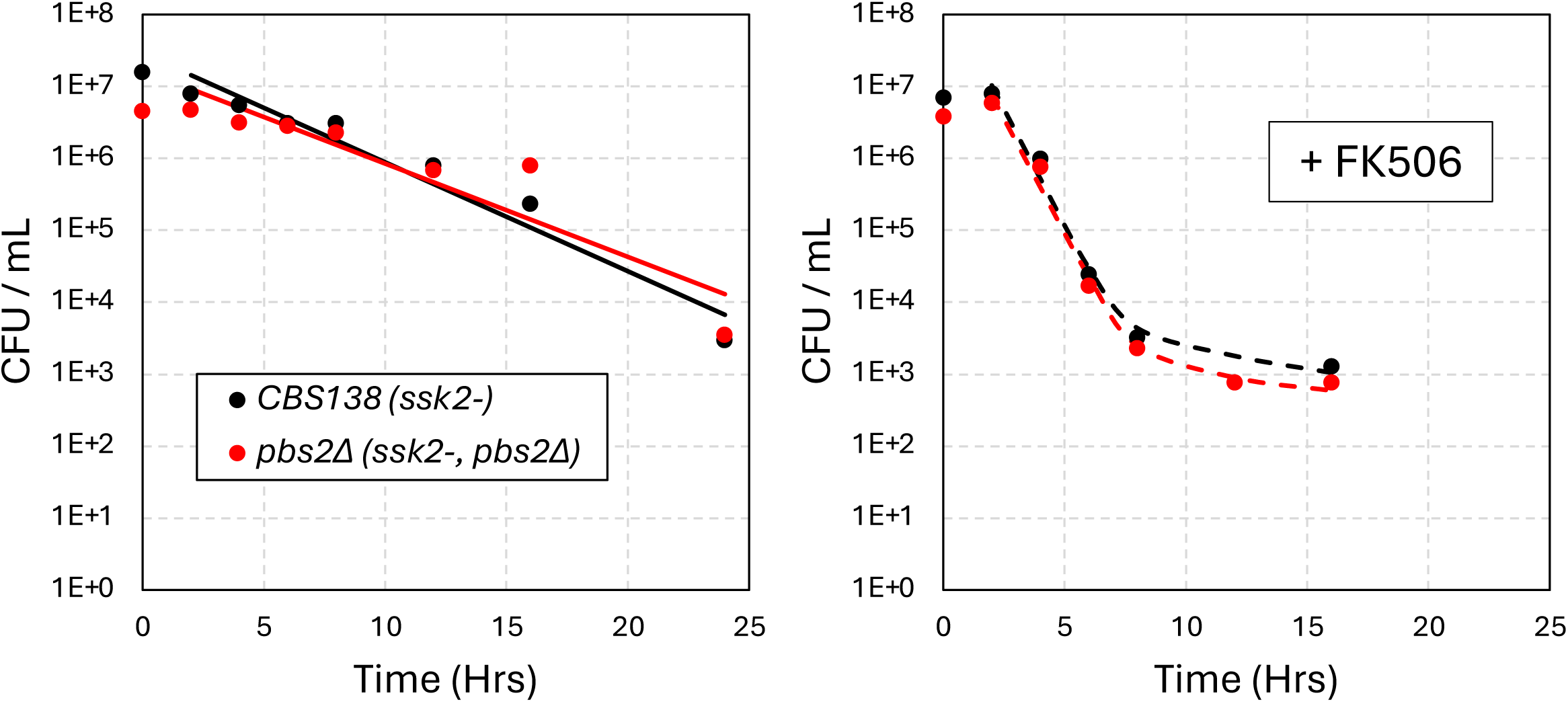
CBS138-derived strains naturally lack HOG signaling. Wild-type CBS138-derived with natural *ssk2-* mutation and a *pbs2Δ* derivative were grown, treated with a high dose of micafungin (1 µg/mL) containing or lacking FK506 (1 µg/mL), and sampled as described in Fig. 1. Data points are the averages of four biological replicates and were used to generate the curve fits.

